# Benchmarking statistical methods for analyzing parent-child dyads in genetic association studies

**DOI:** 10.1101/2021.09.16.460702

**Authors:** Debashree Ray, Candelaria Vergara, Margaret A. Taub, Genevieve Wojcik, Christine Ladd-Acosta, Terri H. Beaty, Priya Duggal

## Abstract

Genetic association studies of child health outcomes often employ family-based designs. One of the most popular family-based designs is the case-parent trio design that considers the smallest possible nuclear family consisting of two parents and their affected child. This trio design is particularly advantageous for studying relatively rare disorders because it is less prone to type 1 error inflation due to population stratification compared to population-based study designs (e.g., case-control studies). However, obtaining genetic data from both parents is difficult, from a practical perspective, and many large studies predominantly measure genetic variants in mother-child dyads. While some statistical methods for analyzing parent-child dyad data (most commonly involving mother-child pairs) exist, it is not clear if they provide the same advantage as trio methods in protecting against population stratification, or if a specific dyad design (e.g., case-mother dyads vs. case-mother/control-mother dyads) is more advantageous. In this article, we review existing statistical methods for analyzing genome-wide data on dyads and perform extensive simulation experiments to benchmark their type I errors and statistical power under different scenarios. We extend our evaluation to existing methods for analyzing a combination of case-parent trios and dyads together. We apply these methods on genotyped and imputed data from multi-ethnic mother-child pairs only, case-parent trios only or combinations of both dyads and trios from the Gene, Environment Association Studies consortium (GENEVA), where each family was ascertained through a child affected by nonsyndromic cleft lip with or without cleft palate. Results from the GENEVA study corroborate the findings from our simulation experiments. Finally, we provide recommendations for using statistical genetic association methods for dyads.

## Introduction

Studies of genetic associations of most human traits and diseases focus on population-based designs (e.g., case-control studies, cohort studies, or data from biobanks) especially for complex and heterogeneous disorders, where both genetic and environmental risk factors are likely involved. For rare disorders this often requires amassing cases sampled from multiple populations, which creates the possibility of type I error inflation due to confounding (as termed by epidemiologists) or population stratification (as referred to by geneticists). Family-based designs, on the other hand, play an important role in the investigation of genetic underpinnings of low frequency or rare disorders (e.g., birth defects) [Benyamin, Visscher and McRae, 2009]. The case-parent trio design is one of the most popular family-based designs which consists of an affected child (i.e., the proband) and both parents. Statistical methods focused on transmission of variants within families, such as in case-parent trio design, protect against population stratification [Schwender et al., 2012]. However, genetic information on both parents is often not available because it is more difficult to recruit biological fathers, which often leads to an abundance of information on mother-child dyads [Shi et al., 2008]. As more multi-ethnic studies become possible, the value of family-based methods that are less prone to population stratification are key for studies of childhood diseases. It is essential to define the best suited analytical approach for each alternative family-based scenario, especially in the presence of a predominance of dyads.

Although statistical methodologies for population-based genome-wide association studies (GWAS) continue to evolve, less ongoing attention has been devoted to methods for family-based studies. While there exist methods for analyzing dyads, it is not clear if one method is consistently more advantageous over another for a given design, and if they have the same robustness against population stratification as full case-parent trios. Furthermore, methods for analyzing dyads were mostly developed 20 or more years ago when type I error performance and statistical power were evaluated at nominal significance levels instead of the more stringent genome-wide levels used now. Recently, Hecker, Laird and Lange [2019] compared type I error and power of methods for general pedigree designs under different founder genotype distribution schemes but only considered the case-parent trio design among nuclear family designs and no hybrid designs. Gjerdevik et al. [2020] compared the relative efficiency of different hybrid designs with the case-control and the case-parent trio designs using direct and indirect effects under log-linear models implemented in the HAPLIN program. However, Gjerdevik et al. [2020] focused on log-linear models only, power/sample size comparisons without any type I error calibration or compute time comparison, and were restricted to a homogenous racial/ethnic group.

In this paper, we benchmark multiple open-source statistical popular methods across both dyad and trio designs on multi-ethnic samples in terms of their compute times, type I error control, and statistical power for identifying common variant associations at stringent significance levels. Using large-scale simulations, we compare and evaluate methods for three nuclear family designs (case-parent trios, case-mother dyads, and a combination of case-parent trios and case-mother dyads), plus two related hybrid designs (case-parent/control-parent trios, case-mother/control-mother dyads) under different parameter settings. Then, we apply these methods to genotyped and imputed data from multi-ethnic mother-child pairs only or trios only or combinations of both dyads and trios from the Gene, Environment Association Studies consortium (GENEVA).

## Material and Methods

### Model and notation

In the following, we consider a GWAS of *N* individuals collected under different nuclear family study designs, particularly case-parent trios and case-parent dyads (typically mother-child pairs). All individuals are genotyped/imputed or sequenced genome-wide at *p* genetic variants (where *p* is of the order of millions), and their disease status of interest is measured. For simplicity, we only consider data on bi-allelic single nucleotide polymorphisms (SNPs). We are interested in testing for association between a SNP and the disease status. In the following subsections, we provide a brief overview of existing methods for analyzing different nuclear family and related hybrid designs.

### Methods for case-parent trios

Consider a study of *n* case-parent trios, where *n* affected (case) offspring are ascertained from a population, and the affected offspring along with their parents are genotyped/imputed or sequenced. Whether information on parents’ disease status is needed depends on the statistical method used. This is the simplest family-based design and existing approaches for analyzing such data includes variations of the transmission disequilibrium test and mating-type-stratified regression approaches.

#### Transmission disequilibrium test (TDT)

The TDT was originally proposed to assess if one allele at a SNP is transmitted from heterozygous parents to their affected offspring more often than that expected under strict Mendelian transmission [Spielman, McGinnis and Ewens, 1993]. This would indicate that the SNP being tested (or marker locus) is both linked and associated with a causal SNP (or disease-susceptibility locus, DSL) for the disease. Essentially, the TDT is a non-parametric test that does not require assumption about disease model or disease distribution in the population [Laird and Lange, 2006]. Note, although the TDT does not require prior specification of any disease model, it implicitly assumes multiplicative effects of alleles [Fallin et al., 2002]. It is also referred to as the allelic TDT, and it boils down to McNemar’s test for a 2×2 contingency table that cannot accommodate covariates or provide estimates of relative risks (RRs). For a large enough sample size, the TDT statistic has a *χ*^2^ distribution with 1 degree of freedom (df) under the null hypothesis of no association *or* no linkage between the marker SNP and an unobserved DSL.

Although the TDT was originally proposed to test for linkage in the presence of association, it is now typically used as a test for association [Laird and Lange, 2006] where the goal is to detect a disease-associated SNP by considering alleles at several markers in a region sequentially rather than a single marker SNP in linkage disequilibrium (LD) with an unobserved DSL [Fallin et al., 2002]. Presence of linkage but no association between the marker SNP and a DSL results in no association between the disease and the marker SNP [Laird and Lange, 2006]. Presence of both linkage and association between the marker SNP and a DSL indicates presence of association between the disease and the marker SNP being tested. Henceforth, any reference to the null hypothesis refers to no association between the disease and the marker SNP.

#### Genotypic TDT (gTDT)

The gTDT compares the case’s transmitted genotype at the SNP to the set of all possible genotypes the case could have inherited from the parental genotypes [Self et al., 1991, Schaid, 1999]. Unlike the TDT, the gTDT considers individuals as units of analysis; can accommodate family-level covariates; assumes some pre-specified genetic inheritance model; and yields estimated RRs with their standard errors [Fallin et al., 2002]. The gTDT uses a conditional logistic regression model and involves a numeric likelihood maximization that can be computationally burdensome at a genome-wide level. However, recently derived closed-form parameter estimates enable rapid genome-wide application of the gTDT when testing additive, dominant or recessive effects using Wald’s test [Schwender et al., 2012]. The gTDT statistic has an asymptotic 1-df *χ*^2^ distribution under the null hypothesis.

#### Generalized disequilibrium test (GDT)

This is a generalization of TDT-like family-based association method proposed to take advantage of larger pedigree information [Chen, Manichaikul and Rich, 2009]. It assesses the genotypic differences between all discordant relative pairs; can model covariates and uses a score test, where the score is obtained from the quasi-likelihood function for a conditional logistic regression model. Under the null hypothesis, the GDT statistic has an asymptotic *N*(0,1)distribution that does not depend on the inheritance model for the DSL. For case-parent trios, a special case of the GDT called the GDT-PO is used where the score is a weighted sum of genotypic differences between phenotypically discordant parent-child pairs.

#### Mating-type-stratified conditional likelihood approach

Similar to genotype RR modeling of Schaid and Sommer [1993], Fan et al. [2013] derived the conditional likelihood of parental mating type and offspring genotype at a SNP given the affected status of offspring under a specific inheritance model assuming Hardy-Weinberg equilibrium (HWE) and random mating in the parental generation. This approach can handle missing parental data without imputation, and can be applied to data on trios, dyads and monads (henceforth, it is referred to as the TDM). The likelihood function is modeled using the two unknown genotypic relative risk (GRR) parameters and the minor allele frequency (MAF) at the marker SNP, obtained using the Newton-Raphson method. Additive, dominant, recessive or multiplicative effects may be assumed, and the resultant likelihood ratio test (LRT) statistic for the TDM has an approximate 1-df *χ*^2^ distribution under the null hypothesis. One need not assume any inheritance model (i.e., assume an unrestricted model) and the resultant TDM statistic has an approximate 2-df *χ*^2^ distribution under the null hypothesis.

#### Log-linear modeling approach

The log-linear approach generalizes the TDT to include orthogonal tests of offspring vs. maternal genetic factors, and can accommodate different risk conferred by a single copy vs. two copies of a risk allele [Weinberg, Wilcox and Lie, 1998]. This approach provides RR estimates for offspring genotype and is generalizable to a wide range of causal scenarios. It lists all possible trio genotypes stratified by parental mating type and applies a Poisson regression to the expected counts of the different trio genotypes conditional on the affected status of the offspring. Inferences about association is carried out using asymptotically *χ*^2^-distributed LRT statistic. The *χ*^2^ df under the null is dictated by the number of GRR parameters, which in turn depends on the inheritance model and the causal scenario assumed. [van den Oord and Vermunt, 2000] described how this log-linear approach can be implemented in LEM, a general computer program for analyzing categorical data. Later, a dedicated genetics software package, HAPLIN, was developed to implement such log-linear models not only for bi-allelic variants but also for multi-allelic variants and other generalizations [Gjessing and Lie, 2006, Gjerdevik et al., 2019]. Using LEM or HAPLIN, one can test for offspring effects only using a 2-df test, maternal effects only in a separate 2-df test or both offspring and maternal effects in a 4-df test. P-values from the 2-df test of offspring effects are directly comparable to p-values from other TDT-like methods.

### Methods for case-mother dyads

In any study of nuclear families, genetic measurements are frequently missing for one parent. More often than not, fathers are missing: they can be harder to recruit, and paternity is inherently harder to be confident of than maternity [Shi et al., 2008]. Consider a study of *n* case-mother dyads, where *n* affected (case) offspring are sampled from the population, and the affected offspring and their mothers are genotyped/imputed or sequenced. One might come up with a straightforward approach of applying the TDT (or any method for case-parent trio data) on such family pairs for whom the genotype of the father at the SNP of interest can be unambiguously inferred. However, this process of selectively including only unambiguous dyads and discarding ambiguous ones can lead to invalid inference due to biases that depend heavily on allele frequencies [Curtis and Sham, 1995].

#### TDT-like approaches

The first appropriate methodological development for the analysis of nuclear family data with missing genetic information on one parent was the 1-TDT [Sun et al., 1999]. It examines the difference between the GRRs of heterozygotes vs. homozygotes (two possible choices) by using all heterozygous parent-homozygous offspring and homozygous parent-heterozygous offspring pairs [Sun et al., 1998, Sun et al., 1999]. Under the null hypothesis, these two GRRs are expected to be the same and the 1-TDT test statistic has an approximate *N*(0,1)distribution. If the total number of afore-mentioned parent-child pairs is not large, an exact p-value can also be calculated under the binomial distribution. The GDT-PO can also be directly applied to data with 1 missing parent. Unlike the 1-TDT, GDT-PO examines all heterozygous parent-homozygous offspring and homozygous parent-heterozygous offspring assuming all available parents are unaffected (recall, GDT only uses phenotypically discordant pairs). For a dataset where all offspring are affected and all their parents are unaffected, 1-TDT and GDT-PO become identical. The GDT-PO statistic has an asymptotic *N*(0,1)distribution under the null hypothesis.

#### Mating-type-stratified likelihood approaches

The TDM is another approach that may be applied to data on parent-child dyads alone and gives a 1-df or a 2-df *χ*^2^ test statistic under the null depending on whether a specific inheritance model is assumed or not. The log-linear approach, too, is flexible enough to handle missing genetic data on parents via the Expectation-Maximization (EM) algorithm [Weinberg, 1999], and can be implemented using programs such as LEM or HAPLIN.

### Methods for case-mother dyads and case-parent trios combined

When conducting family studies, it is not always possible to collect only families of one structure. In practice, we may not have only case-parent trios or only case-mother dyads but combinations of both. Suppose we have *n*_1_ complete case-parent trios and *n*_2_ incomplete trios where, without loss of generality, the fathers are missing. Note, it does not matter here if the fathers or the mothers are missing since we are only assessing direct effects of inherited genotypes of offspring on their disease status.

#### TDT-like approaches

Instead of ignoring one set of families depending on whether the sample size *n*_1_ is larger than *n*_2_ or not, [Sun et al., 1999] proposed applying the TDT on all complete trios and the 1-TDT on all dyads and then combining the two statistics. The resultant combined statistic, denoted TDT_com_, has an asymptotic *N*(0,1)distribution. However, there is currently no software for TDT_com_. One can instead implement the 1-TDT or the GDT-PO on all parent-child pairs without discarding any families.

#### Mating-type-stratified likelihood approaches

As described before, the TDM and the log-linear models can also be used in this scenario.

### Methods for case-parent/control-parent trios

Genetic association studies, whether family-based (e.g., case-parent trio design) or population-based (e.g., case-control design), have their own strengths and limitations; see Weinberg and Umbach [2005] for a comprehensive summary of their advantages and disadvantages. A hybrid design bringing the strengths of case-parent trio and case-control designs into a single analytic framework is the case-parent/control-parent trio design. Consider a study of *n* trios, where *n*_1_ affected (case) and *n*_2_ = *n* − *n*_1_ unrelated unaffected (control) offspring are sampled from the population, and all sampled offspring and their parents are genotyped/imputed or sequenced. Although control-parent trios are generally easier to recruit and, along with case-parent trios, can help guard against spurious signals due to segregation distortion, they are typically either not recruited or discarded from analysis even when available because the TDT or the gTDT is only applicable to trios with affected offspring [Deng and Chen, 2001].

#### TDT-like approaches

A straightforward approach is to apply the TDT on case-parent trios alone (refer to this as TDT_D_) and on control-parent trios separately (refer to this as TDT_C_, which, in contrast to TDT_D_, can be viewed as a test of transmission of the ‘non-risk’ allele at the DSL rather than the ‘risk’ allele), and then combine these two independent tests into a new test TDT_D+C_ [Deng and Chen, 2001]. This TDT_D+C_ statistic has an asymptotic 2-df *χ*^2^ distribution under the null hypothesis. [Deng and Chen, 2001] additionally proposed TDT_DC_, a contingency table association test of allele transmissions (from heterozygous parents) with disease status in unrelated offspring. This TDT_DC_ statistic has an approximate 1-df *χ*^*2*^ distribution under the null hypothesis and does not require equal numbers of case-parent and control-parent trios.

#### Mating-type-stratified likelihood approaches

A log-linear model can be used to combine the family-based case-parents-trio component and the population-based parent-parent component, thus not requiring genetic data on the control offspring [Weinberg and Umbach, 2005]. It assumes the disease is rare for the offspring of each parental genotype combination, mating symmetry, and Mendelian proportions in the population. It neither assumes Hardy-Weinberg equilibrium nor random mating. In the presence of population structure, one can generalize this log-linear model to include a disease-status by total-number-of-parental-alleles interaction term or 5 additional disease-status by mating-type interaction terms (note, 6 distinct unordered parental mating types are possible here). However, “[a] direct consequence of preferring the enlarged model is that the control portion of the data will not contribute to inference related to the risk parameters, and the population-based component, in effect, becomes statistically irrelevant” [Weinberg and Umbach, 2005].

### Methods for case-mother/control-mother dyads

Consider a study of *n* dyads, where *n*_1_ affected (case) offspring are ascertained, and *n*_2_ = *n* − *n*_1_ unaffected (control) offspring are sampled. Genotype data are available on offspring and their mothers. There are fewer methods for this hybrid dyad design. No TDT-like methods have been proposed for this design but the log-linear approach using LEM or HAPLIN is applicable [Shi et al., 2008].

### Simulation experiments

We first simulate 1,000 trios using the LE program [Chen and Deng, 2001, Chen, Manichaikul and Rich, 2009]. For the relevant scenarios detailed below, we remove fathers to obtain mother-child pairs (dyads). The LE program is a general program for simulating pedigrees requiring the following parameters as input: disease prevalence, disease allele frequency, genotypic penetrances, number of families, and structure of the pedigree. We simulate only one causal SNP at a time with a fixed MAF to ensure independence of SNPs (which is needed to estimate type I error rate and statistical power). We simulate only offspring GRR effects and assume different GRR values to generate both null and non-null SNPs. For our type I error and power analyses, we simulate 1 million null SNPs (GRR=1) and 10,000 non-null SNPs, respectively. For non-null SNPs, we assume GRR=2 under two different inheritance models (additive and multiplicative). Note, for GRR=1, the inheritance model does not influence the values of genetic penetrance. Our choices of these parameters are described below for specific scenarios.

We evaluate two classes of methods: TDT-like methods (TDT, gTDT, 1-TDT, GDT-PO, TDT_DC_, TDT_D+C_) and the mating-type-stratified likelihood approaches (HAPLIN). While the TDM and log-linear modeling using LEM are also candidates for mating-type-stratified likelihood approaches, we exclude both. In many analyses, TDM faced issues with matrix inversion that prevented successful model fit, leading to missing results for many SNPs. We also faced several roadblocks in implementing current LEM executable genome-wide on a Unix cluster. Note, not all methods in each class are applicable for a given study design. We simulate two scenarios involving samples from either one or two homogenous ancestral populations mimicking situations without or with population stratification. While we benchmark type I error of methods for all simulation settings, only statistical power for the homogenous group is used for benchmarking since not all methods could maintain type I error in the presence of population stratification. We use QQ plots to evaluate type I error rates at stringent levels, and compare power calculated at conventional genome-wide level (5 × 10^−8^).

#### Scenario 1: One homogenous genetic ancestry

All the parents are simulated from a single homogenous ancestral population with disease prevalence of 30%. A fixed MAF of 10% is assumed for the causal SNP. Under this scenario, both type I error and power are compared for all methods.

#### Scenario 1A: Case-mother dyads

The fathers from 1,000 case-parent trios are removed to obtain 1,000 case-mother dyads.

#### Scenario 1B: Case-mother dyads and case-parent trios combined

We remove fathers from the first 750 case-parent trios to obtain 750 case-mother dyads (75% of the dataset), leaving the remaining 250 case-parent trios (25% of the dataset).

#### Scenario 1C: Case-mother/control-mother dyads

We generate 500 case-parent trios and 500 control-parent trios independently. Removing the father from each trio results in 500 case-mother dyads and 500 control-mother dyads, which are analyzed together. For power calculations, besides this 50:50 case-control ratio among offspring, we also check 70:30 and 30:70 ratios of case-control families.

#### Scenario 2: Two distinct genetic ancestry groups

This scenario considers existence of population substructure between families in the sample. We simulate the parents of 500 families from one homogenous ancestral population with disease prevalence of 30%, and the parents of the other 500 families from a separate ancestral population with a lower disease prevalence of 15%. The causal SNP is simulated to have an MAF of 10% and 3% respectively in the two populations. We analyze a pooled sample for all methods. Additionally, for the mating-type-stratified likelihood approach, we apply HAPLIN to each ancestry group separately and then meta-analyze using Fisher’s p-value combination method [Fisher, 1925, Ray, Pankow and Basu, 2016]. Under this scenario, we compare type I error rates only.

#### Scenario 2A: Case-mother dyads

Fathers from all 1,000 case-parent trios (500 from each ancestral population) are removed to obtain 1,000 case-mother dyads.

#### Scenario 2B: Case-mother dyads and case-parent trios combined

From each ancestral group, we remove fathers of the first 75% of case-parent trios to obtain 750 case-mother dyads in total and combined this with the remaining 250 case-parent trios. This resulted in 375 case-parent trios and 125 case-mother dyads from each ancestral population.

#### Scenario 2C: Case-mother/control-mother dyads

We generate 500 case-parent trios and 500 control-parent trios independently. Among case-parent trios, 250 are simulated for each ancestral population. Similarly for the control-parent trios. Removing the father from each trio gives 500 case-mother and 500 control-mother dyads in a genetically heterogeneous sample.

### Application to GENEVA data on orofacial clefts

In GENEVA, case-parent trios were ascertained through cases with an isolated, nonsyndromic orofacial cleft (i.e., cleft lip; cleft palate; or cleft lip with palate). They were largely recruited through surgical treatment centers by multiple investigators from Europe (Norway and Denmark), the United States (Iowa, Maryland, Pennsylvania, and Utah) and Asia (People’s Republic of China, Taiwan, South Korea, Singapore, and the Philippines) over several years [Beaty et al., 2010]. Type of cleft, sex, race, family history, and common environmental risk factors were collected through direct maternal interview. Genotyping on the Illumina Human610 Quadv1_B array with 589,945 SNPs was performed at the Center for Inherited Disease Research (https://cidr.jhmi.edu/). As part of two recent publications [Zhang et al., 2021, Ray et al., 2021], trio-aware phasing and re-imputation using the 1000 Genomes Phase 3 release 5 reference panel were performed. Among GENEVA participants who were re-imputed and used in these two articles, we restrict our analysis to the participants ascertained through nonsyndromic CL/P. We use ‘hard’ genotype calls: if the calls had uncertainty >0.1 (i.e., genotype likelihoods <0.9), they are treated as missing; the rest are regarded as observed genotype calls. All imputed SNPs were filtered to exclude any with *R*^2^<0.3. All variants are on the forward strand.

For the current analysis, we analyze genotyped/imputed SNPs only and take the following quality control measures using PLINK 1.9 [Chang et al., 2015]: all SNPs with MAF<5% and any showing deviation from Hardy-Weinberg equilibrium (HWE) at *p* <10^−6^ among parents are excluded; all genotyped SNPs with missingness >5% and Mendelian error rate >5% are also removed. Additionally, all trios with per-trio Mendelian error rate >5% are dropped. Our final GENEVA analytical dataset contains 5,204,784 autosomal SNPs, including both observed and imputed SNPs having MAF >5% among parents, for 1,487 multi-ethnic complete case-parent trios. Of these 1,487 complete trios, 891 trios are of Asian ancestry (including Malays from Singapore) and 575 are of European ancestry, respectively. The remaining 22 trios are from other racial/ethnic groups. Among 2,974 parents, 560 have missing phenotype information. There are 534 female and 953 male CL/P probands in total.

We analyze three separate designs: case-parent trios only (considered here as the gold standard); case-mother dyads only; and trios and dyads combined. For the trios only dataset, we analyze all 1,487 case-parent trios (a multi-ethnic sample). For the dyads only dataset, we remove the father from each trio and analyze the resulting 1,487 case-mother pairs. For the combined dataset, we remove the father from 75% of all trios within each racial/ethnic group (consistent with the simulations for scenario 2B detailed above), yielding 1,116 multi-ethnic case-mother pairs and 371 multi-ethnic case-parent trios. We compare findings from the dyads only and the combined designs against the trios only design. Note, there are no control families in GENEVA, and hence no case-parent/control-parent or case-mother/control-mother design are considered. We analyze the complete GENEVA dataset using TDT-like methods. We apply HAPLIN to the Asian and the European groups separately and then meta-analyze results from the 2-df test for offspring genotype effects using Fisher’s p-value combination. We exclude the 4-df combined test of offspring or maternal effects in this comparison since we cannot rule out the influence of maternal genes on risk of clefts in offspring [Jugessur et al., 2010, Shi et al., 2012].

For each analysis, we define independent loci by clumping all the genome-wide significant SNPs (*p* < 5× 10^−8^) in a ±500 Kb span and with LD *r*^2^ > 0.2 into a single genetic locus. We used the SNP2GENE function of FUMA (v1.3.6b) [Watanabe et al., 2017] for clumping and mapping each locus to the gene nearest to the lead SNP. The index SNP for each locus is chosen as the most significant SNP. Since we perform multi-ethnic analysis, we separately use 1000G Phase 3 EUR and EAS as reference populations for LD calculation. For a given analysis, we find independent hits to be the same regardless of the ancestry of the reference group chosen for LD calculation. We define the bounds of a locus as the minimum of lower bounds and the maximum of upper bounds across both ancestry groups. All genomic coordinates are given in NCBI Build 37/UCSC hg19.

## Results

### Simulation experiments: Type I error

#### Scenario 1A: One homogenous ancestry group, case-mother dyads

The QQ plots of all methods, except 1-TDT and GDT-PO, appear to fully lie within the 95% confidence interval (CI) for the expected distribution of p-values, thus indicating their correct type I error control (**Figure 1**). The 1-TDT and GDT-PO perform well at nominal error levels (*λ*_1−*TDT*_ =1 and *λ*_*GDT*−*PO*_ =1) but show some inflation at stringent error levels as indicated by the data points falling outside the 95% CI. The corresponding gold standards for all methods (i.e., methods applied to case-parent trios) maintain type I error as well. As expected, when all offspring are affected and all parents are unaffected, the 1-TDT and the GDT-PO curves coincide perfectly.

**Figure 1:**
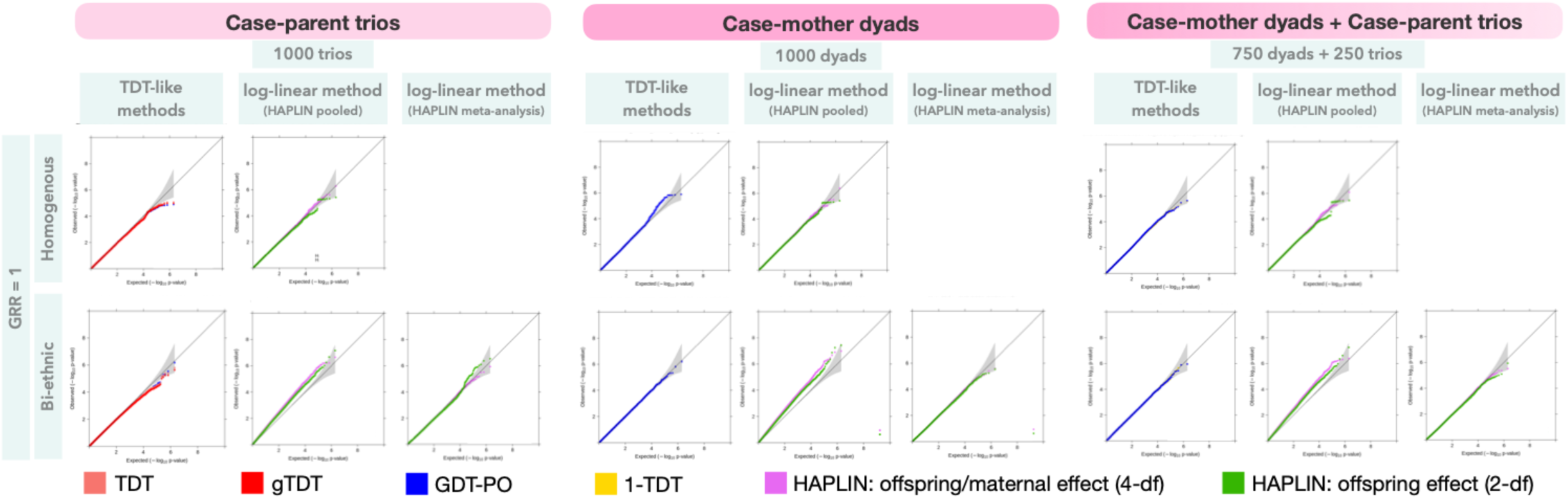
Type I error performance of the different combinations of methods and nuclear-family designs at stringent significance levels. Results are based on simulated data on either one or two homogenous racial/ethnic groups with 1 million null SNPs. For one homogenous sample, a common disease prevalence of 30% and MAF 10% was simulated. For bi-ethnic data, the second homogenous group had a disease prevalence of 15% and MAF 3%. All offspring were affected, and all parents were unaffected. Observed(−log_10_p-values) are plotted on the y-axis and Expected(−log_10_p-values) on the x-axis of these QQ plots. The gray shaded region in each QQ plot represents a conservative 95% confidence interval for the expected distribution of p-values.

#### Scenario 1B: One homogenous ancestry group, case-mother dyads and case-parent trios combined

All methods perform similar to the corresponding gold standard methods, maintaining appropriate type I error even at stringent levels (**Figure 1**).

#### Scenario 1C: One homogenous ancestry group, case-mother/control-mother dyads

HAPLIN, the only applicable method in this scenario, show well-controlled type I error (**Figure 2**). Note, for the corresponding case-parent/control-parent trio data, both HAPLIN and the TDT-like methods maintain correct type I error at stringent levels.

**Figure 2:**
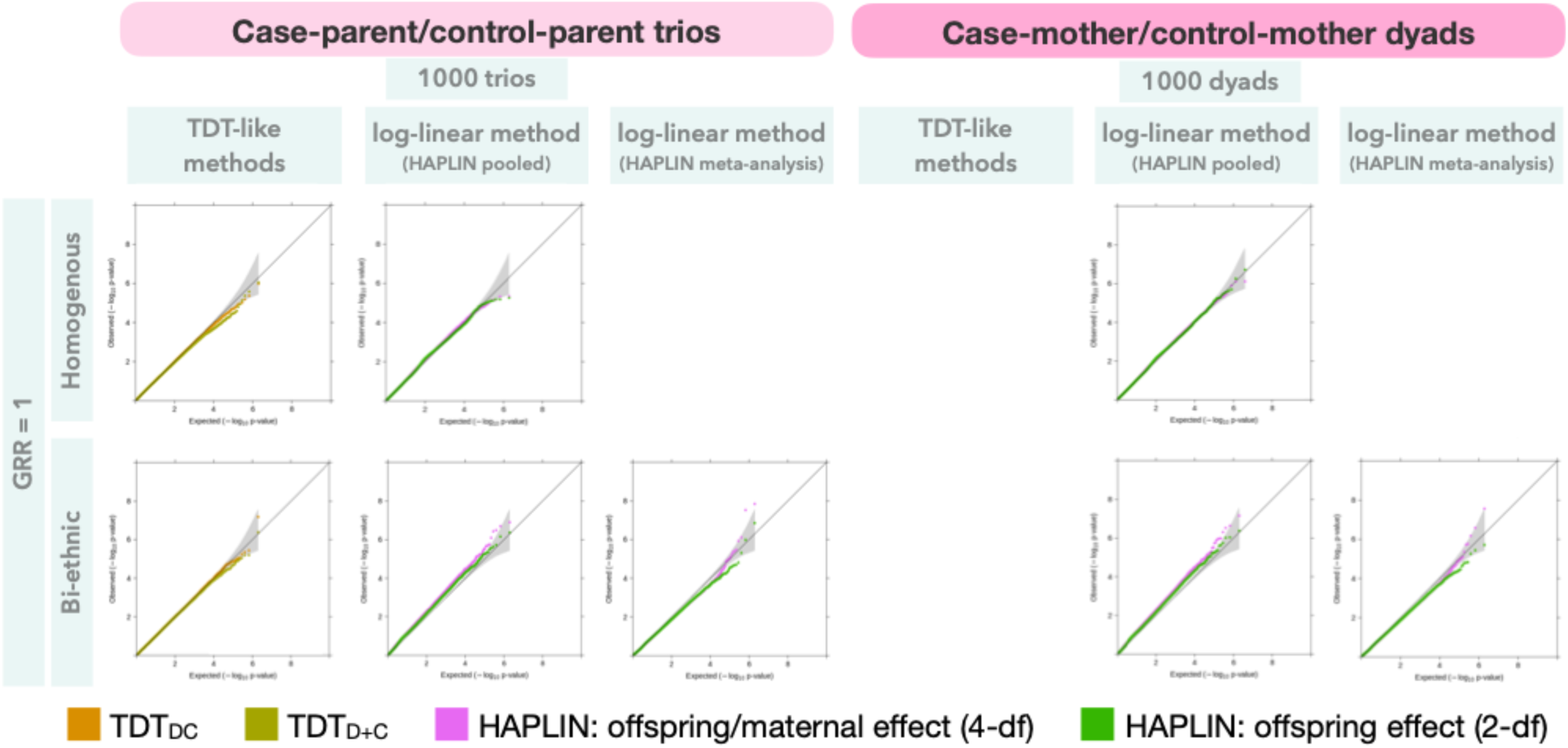
Type I error performance of the different combinations of methods and hybrid family designs at stringent significance levels. Results are based on simulated data on 1,000 families from either one or two homogenous racial/ethnic groups with 1 million null SNPs. For one homogenous group, a common disease prevalence of 30% and MAF 10% was simulated. For bi-ethnic data, the second homogenous group had a disease prevalence of 15% and MAF 3% for the causal SNPs (note, null effect is assumed). Case-to-control ratio among offspring was 50:50, and all parents were unaffected. Observed(−log_10_p-values) are plotted on the y-axis and Expected(−log_10_p-values) on the x-axis of these QQ plots. The gray shaded region in each QQ plot represents a conservative 95% confidence interval for the expected distribution of p-values.

#### Scenario 2A: Two distinct ancestry groups, case-mother dyads

All TDT-like methods maintain appropriate type I error as indicated by their QQ plots lying within the 95% CI for the expected distribution of p-values (**Figure 1**). HAPLIN show inflated type I error even at nominal significance levels (*λ*_*HAPLIN*−2*df*_ = 1.34 and *λ*_*HAPLIN* −4*df*_ = 1.60). These observations are reflected for the gold standards as well. When HAPLIN is applied to each ancestry group separately and then meta-analyzed, type I error is then well-controlled at nominal (*λ*_*HAPLIN*−2*df*_ = 1.02 and *λ*_*HAPLIN* −4*df*_ = 1.00) as well as stringent levels. However, for the corresponding gold standard, the QQ plots indicate type I error is well-controlled for the 4-df test but is inflated at stringent levels for the 2-df test.

#### Scenario 2B: Two distinct ancestry groups, case-mother dyads and case-parent trios combined

As before, all TDT-like methods exhibit well-controlled type I error rate while HAPLIN show significant type I error inflation (**Figure 1**). Meta-analysis of HAPLIN applied to ancestry-stratified data results in well-controlled type I error for both tests. It is worth noting with real datasets one may not have clearly distinct groups to stratify and then meta-analyze. We also simulate a skewed distribution of dyads and trios within each ancestral group; results indicate robustness of TDT-like methods to population substructure while HAPLIN shows even greater inflation (**Figure S1**).

#### Scenario 2C: Two distinct ancestry groups, case-mother/control-mother dyads

HAPLIN is the only method that could be evaluated here, and it shows considerable type I error inflation when applied to the pooled sample (**Figure 2**). Meta-analysis of HAPLIN applied to ancestry-stratified data results in well-controlled type I error; no different from the corresponding gold standard. Note, we also simulate a skewed distribution of case and control offspring within each ancestral group, with all 500 case families coming from one group and all 500 control families from the other. Our results indicate robustness of TDT-like methods while HAPLIN shows extreme type I error inflation (**Figure S2**).

### Simulation experiments: Power

#### Scenario 1A: One homogenous ancestry group, case-mother dyads

All the TDT-like methods have similar statistical power to detect phenotype-genotype association (**Figure 3a**). HAPLIN is more powerful and, depending on the inheritance model, the 4-df test of offspring/maternal effects show improved power over the 2-df test of offspring effects alone. All methods have reduced power for case-mother dyads compared to those for case-parent trios, as expected.

**Figure 3:**
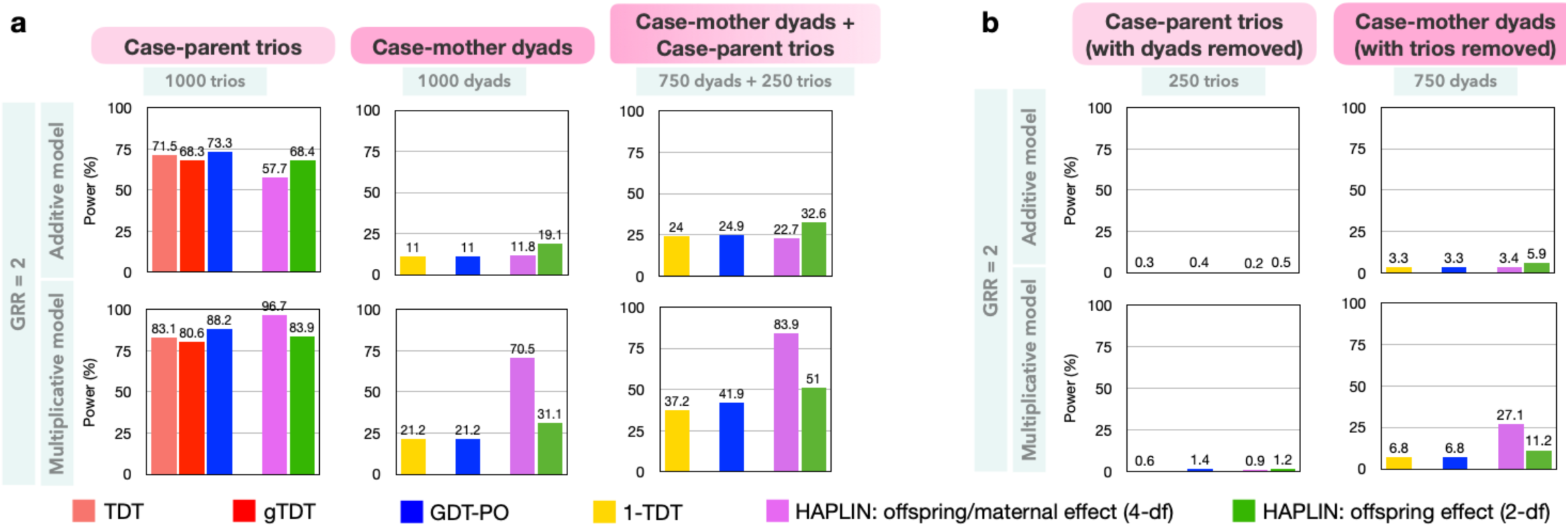
Statistical power for the different combinations of methods and nuclear-family designs at genome-wide significance level (5×10^−8^). Results are based on simulated data on 1,000 families from one homogenous racial/ethnic group with 10,000 non-null SNPs at MAF 10% at the casual SNP, and a common disease prevalence of 30%. All offspring were affected, and all parents were unaffected.**a**. Comparison of designs with the same number of families of different compositions. **b**. Comparison of the combined analysis of 750 case-mother dyads and 250 case-parent trios against the scenarios when either all dyads or all trios are removed from analysis.

#### Scenario 1B: One homogenous ancestry group, case-mother dyads and case-parent trios combined

For a fixed number of families, all methods have improved power over analyzing case-mother dyads alone (**Figure 3a**), as expected. Further, discarding any type of family from a mixed family design like this results in substantial loss of statistical power (**Figure 3b**).

#### Scenario 1C: One homogenous ancestry group, case-mother/control-mother dyads

HAPLIN, the only applicable method, is nearly as powerful in the dyad design as in the full trio design (**Figure 4**). As the proportion of cases among offspring decreases, HAPLIN’s power also decreases. The 2-df test for offspring genotypic effect may or may not be more powerful than the 4-df test of offspring/maternal effect depending on the inheritance model. For the corresponding gold-standard design (i.e., case-parent/control-parent design), there are competing methods although they provide less power than HAPLIN. Between the two TDT-like methods, TDT_D+C_ appear to be at least as powerful as TDT_DC_ regardless of the inheritance model and the case-control ratio among offspring, despite the higher df of TDT_D+C_ (**Figure 4**).

**Figure 4:**
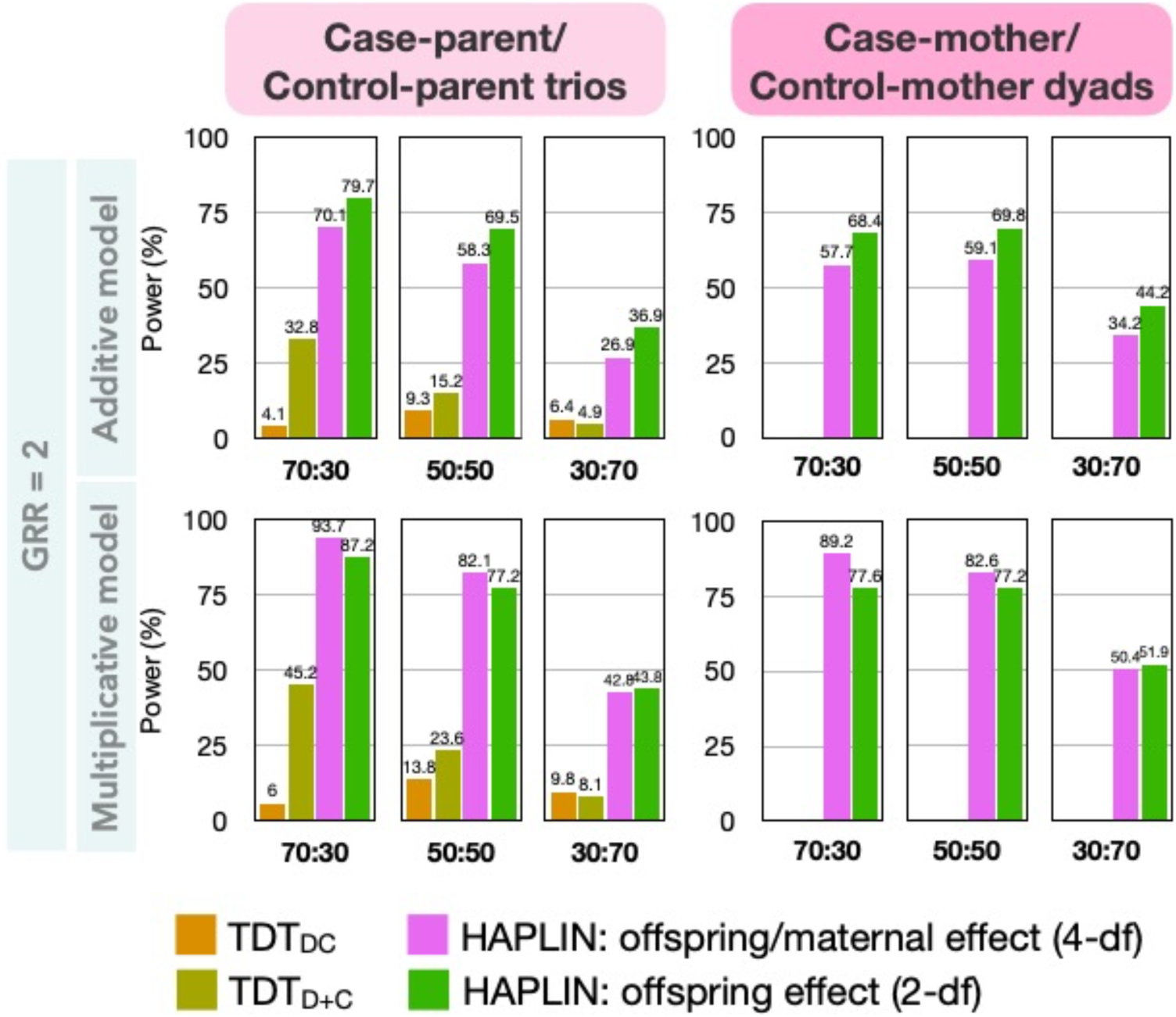
Statistical power for different combinations of methods and hybrid family designs at genome-wide significance level (5×10^−8^). Results were based on simulated data on 1,000 families from one homogenous racial/ethnic group with 10,000 non-null SNPs at MAF 10% at the causal SNP, and a common disease prevalence of 30%. Case-to-control ratio among offspring is either 70:30, 50:50 or 30:70. All parents are unaffected.

### Application to GENEVA Data on Orofacial Clefts

#### Case-parent trios

To establish a gold standard, we first analyze the complete case-parent trio data (pooled sample) using several TDT-like methods (TDT, gTDT and GDT-PO). As expected, TDT and gTDT give nearly identical results with both replicating known signals for CL/P [Dixon et al., 2011, Beaty, Marazita and Leslie, 2016, Ray et al., 2021] at the conventional genome-wide threshold (p<5 × 10^−8^): 1p22.1 (*ABCA4/ARHGAP29*), 1q32.2 (*IRF6*), 8q24 (gene desert), 17p13.1 (*NTN1*), 18q12.1 (*TTR*, not a known cleft-associated region and could be spurious), and 20q12 (*MAFB*) (**Figure 5**). Further, 3p11.1 (*EPHA3*), 8q21.3 (*DCAF4L2*), and 10q25.3 (*SHTN1*) yield suggestive significance (p<10^−6^). The GDT-PO signals are somewhat attenuated compared to those from TDT/gTDT, and it fails to replicate the signals at/near genes *TTR, DCAF4L2* and *SHTN1*. This is likely due to the reduced sample sizes from considering only phenotypically discordant parent-child pairs since some parents had more subtle ‘microforms’ or missing phenotype information.

**Figure 5:**
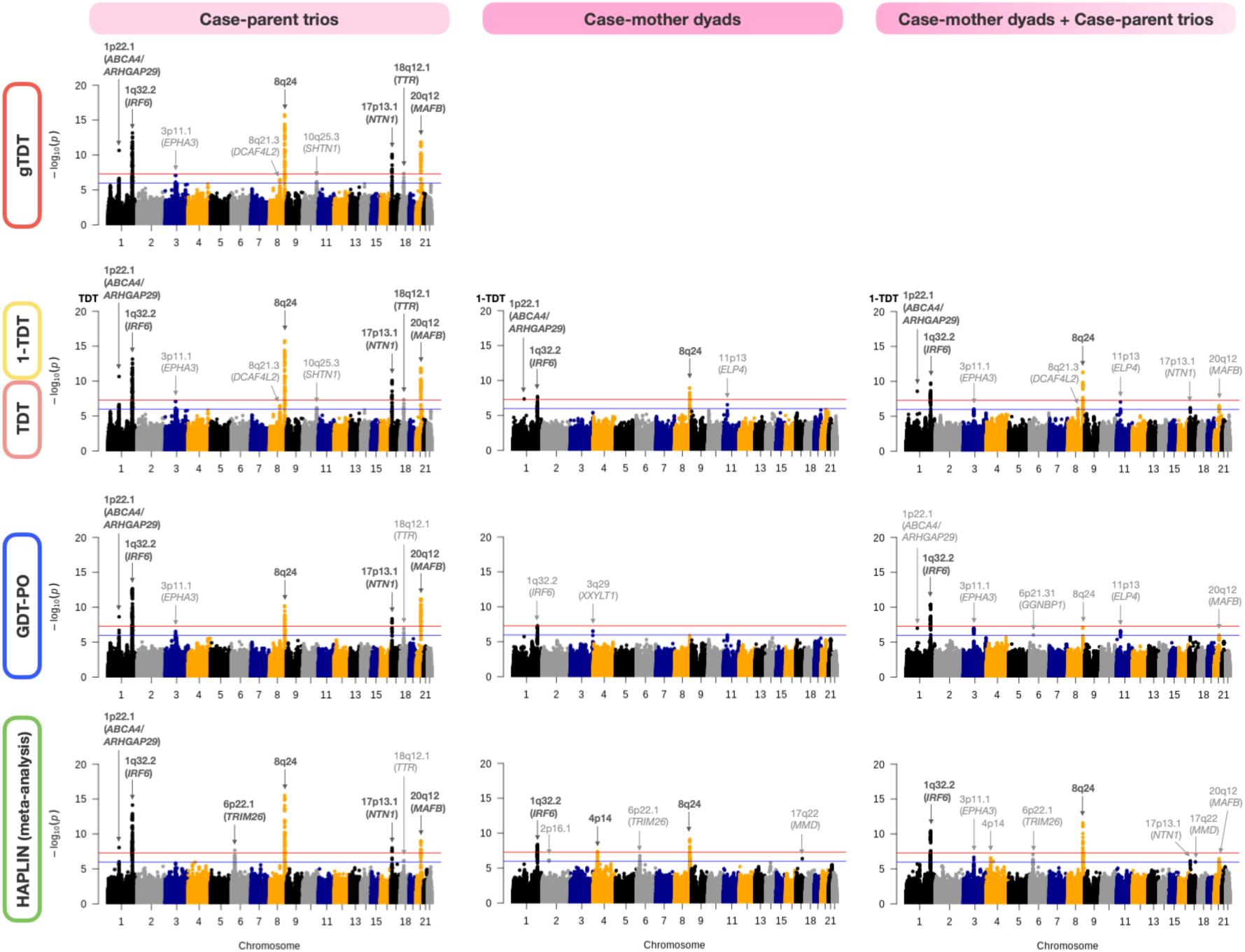
Manhattan plots for the different combinations of methods and nuclear family designs from the multi-ethnic GENEVA study on CL/P. The gTDT and the TDT are applicable to case-parent trio design (*χ* =1487) only. The 1-TDT (a generalization of TDT) is applicable to both case-mother dyad design (*χ* =1487) and the combined case-mother dyad case-parent trio design (*n*_1_=371 trios, *n*_2_ =1116 dyads). The GDT-PO and HAPLIN methods are applicable to all three designs. Here, HAPLIN (2-df test of offspring genotypic effect) was applied on each racial/ethnic group separately and then meta-analyzed. The red and blue horizontal lines in each plot correspond to genome-wide (5×10^−8^) and suggestive (10^−6^) significance levels, respectively. The genome-wide significant loci for each method-design pair are annotated in dark gray and the suggestively significant loci in light gray.

We additionally analyze the complete trios using HAPLIN. Unlike TDT-like methods, HAPLIN is not immune to type I error inflation due to population stratification (**Figure S3**). So, we analyze Asian and European groups separately using HAPLIN and then meta-analyze the results from the 2-df test for offspring effects (**Figure 5**). HAPLIN detects a new signal at 6p22.1 (*TRIM26, p*= 2*R*1 × 10^−8^), which has no known relevance to CL/P and could be spurious. It fails to detect the *EPHA3* signal.

#### Case-mother dyads

The 1-TDT and GDT-PO show considerably reduced power to detect genetic associations compared to the case-parent trio analysis, as expected (**Figure 5**). At genome-wide significance, 1-TDT identifies only the *ABCA4/ARHGAP29, IRF6* and 8q24 signals compared to TDT/gTDT on complete trios. GDT-PO fails to identify any genome-wide significant signals. This lack of power for GDT-PO compared to 1-TDT is presumably in part due to smaller sample sizes and missing phenotype information in some mothers. HAPLIN, when meta-analyzed over Asian and European groups, detects only the *IRF6* and 8q24 signals. It also detects an intergenic region at 4p14 (*p* = 4.2 × 10^−8^), which may be spurious as it is not detected by the HAPLIN meta-analysis of complete trios and has not been previously reported in the cleft literature. It is possible our HAPLIN meta-analysis results may still reflect inflation due to population stratification because each racial/ethnic group is not completely homogenous (participants were sampled from various countries). At the suggestive significance level, a few additional regions are detected, some of which may be spurious since they are not found in the corresponding analysis of the complete trios: 11p13 by 1-TDT; *IRF6* and 3q29 by GDT-PO; 2p16.1, 6p22.1 and 17q22 by HAPLIN.

#### Case-mother dyads and case-parent trios combined

The 1-TDT and GDT-PO show improved power over analyzing the same number of families consisting of case-mother dyads alone (**Figure 5**). At the genome-wide significance level, 1-TDT identifies only the *ABCA4/ARHGAP29, IRF6* and 8q24 signals while GDT-PO identifies only the *IRF6* signal when compared to TDT/gTDT signals for complete trios. HAPLIN, when meta-analyzed over Asian and European groups, detects only the *IRF6* and 8q24 signals. Interestingly, it does not detect the possibly spurious signal at 4p14 at the genome-wide threshold that it identifies from case-mother dyads alone. At the suggestive significance level, 1-TDT detects the *EPHA3, DCAF4L2*, 11p13 (possibly spurious), *NTN1* and *MAFB* signals; the GDT-PO detects the *ABCA4/ARHGAP29, EPHA3*, 6p21.31 (possibly spurious), 8q24, 11p13 (possibly spurious) and *MAFB* signals; and finally, HAPLIN detects the *EPHA3*, 4p14 (possibly spurious), 6p22.1 (possibly spurious), *NTN1*, 17q22 (possibly spurious) and the *MAFB* signals.

### Comparison of Compute Times

We use genetic data on 8,015 SNPs in the region 8q24:128344410-132105518, which includes the known gene desert region. **Figure 6** shows the compute times on an Oracle Virtual Machine 6.1.24 (64bit) with Intel® Xeon® CPU E3-1270 v6 @3.80 GHz processor and 8GB of RAM. Across all three designs, the TDT-like methods have comparable compute times of <1 minute, which include loading and necessary data formatting. HAPLIN applied on the pooled sample takes at least 90 minutes to run for the case-parent trio design and takes 2-3-fold more time for study designs with at most one missing parent. For multi-ethnic samples, application of HAPLIN on ancestry-stratified data and subsequent meta-analysis using Fisher’s method required nearly 1.5-fold increased time compared to HAPLIN applied on the pooled sample.

**Figure 6:**
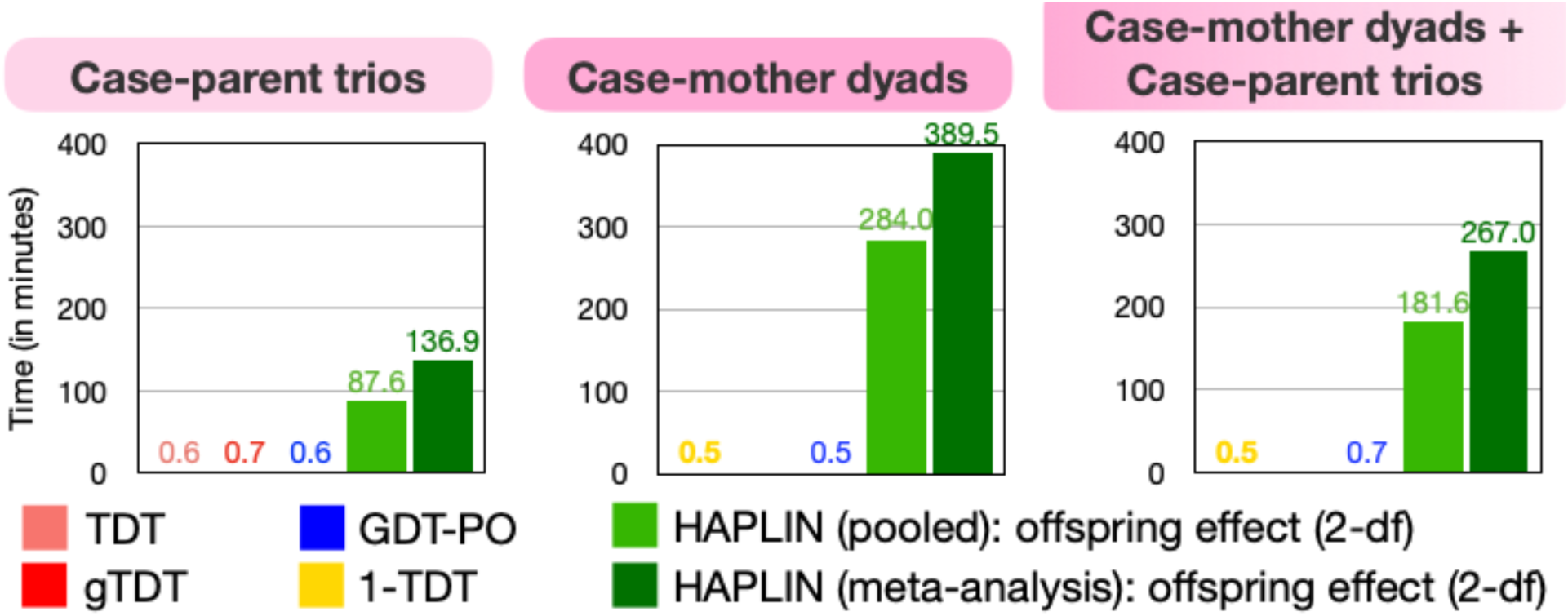
Compute times for the different combinations of methods and nuclear family designs from the multi-ethnic GENEVA study on CL/P. Results are based on a subset of the genetic data: 8,015 genotyped/imputed SNPs in the region chr8:128344410-132105518 that includes the known cleft locus 8q24.

## Discussion

In this article, we review and benchmark existing open-source statistical methods for the genome-wide analysis of parent-child dyads against those available for case-parent trios. We consider case-mother dyads alone, a combination of case-mother dyads and case-parent trios, and combinations of case-mother/control-mother dyads. We compare these study designs against their corresponding gold standards: either case-parent trios alone or case-parent/control-parent trios. We use extensive simulation experiments, and array-based genetic data on trios ascertained through a child affected by CL/P from the GENEVA study. This work is partly motivated by the Environmental influences on Child Health Outcomes (ECHO) study, a cohort collaboration that seeks to identify environmental and genetic exposures relevant to child development and disease. Genetic array data will be paired with a catalog of early childhood or maternal outcomes on more than 30,000 individuals in the United States comprising mother-child-father trios and mother-child pairs across these multi-ethnic ECHO cohorts. Given the expected genetic diversity of these cohorts, it is imperative to evaluate methods that can be applied individually to dyads or trios or in combination while taking maximal advantage of the family-based structure and information whenever possible. Similarly, it is essential to define the analytical approach best suited for each family-based design, especially since dyads are easier to recruit. Recommendations from this study will not only help identify the best strategies to use in diverse studies like the ECHO study, but also provide guidelines for other dyad design-based studies.

### Recommendations

For case-parent trios, the TDT-like approaches (e.g., TDT, gTDT, GDT-PO) inherently circumvent the confounding due to population stratification by distinguishing between transmitted and non-transmitted parental alleles either in a contingency table framework or in a conditional regression framework. The non-transmitted parental alleles in a case-parent trio design– also referred to as the case-parental control design [Sun et al., 1998]– serve as matched genetic controls even when random mating and HWE assumptions are not met [Weinberg, Wilcox and Lie, 1998]. Consequently, generalizations of these methods to accommodate a missing parent in the case-mother dyad design (e.g., 1-TDT, GDT-PO) also protect against confounding due to population stratification.

For multi-ancestry data consisting of either case-mother dyads alone or a combination of case-mother dyads and case-parent trios, both 1-TDT and GDT-PO are useful. We recommend GDT-PO since it can model covariates in a regression framework. If there are many affected parents or parents with missing phenotypes, we recommend 1-TDT since, in this scenario, it is more powerful than GDT-PO, which considers only phenotypically discordant relative pairs. If the dataset consists of a homogeneous genetic ancestry group, we recommend a log-linear approach (e.g., HAPLIN) as it is often more powerful than 1-TDT and GDT-PO. In particular, the 2-df test for offspring genotype effects alone under a log-linear model using LRT tends to be more powerful than the 1-df TDT under a dominant or a recessive genetic model since LRT uses information about the joint transmission from pairs of parents, rather than accounting for the parental transmissions of individual alleles [Weinberg, Wilcox and Lie, 1998, Weinberg, 1999]. If the dataset is multi-ancestry but consists of identifiable genetic ancestry groups, we recommend using a log-linear approach on each homogeneous sub-group and then meta-analyzing the results. However, caution should be exercised in interpreting findings from these log-linear approaches on multi-ethnic data since it is often impossible to ensure each racial/ethnic group in a real dataset is truly homogenous. It is important to highlight that we do not include covariates to adjust for heterogeneity within each group or in the analysis of pooled sample.

For multi-ancestry data consisting of case-parent/control-parent trios, either TDT_DC_ or TDT_D+C_ may be used. Both these methods control type I error even when case- and control-parent trios come from different genetic ancestral populations. Unfortunately, we do not have any recommendation if the data consist of multi-ancestry case-mother/control-mother dyads. A meta-analysis of results from a log-linear approach like HAPLIN can be used if the case-mother/control-mother dyads (or the case-parent/control-parent trios) come from one or more identifiable homogenous genetic ancestry groups. In this case, HAPLIN is usually considerably more powerful than TDT-like approaches; however, few populations are truly homogeneous and for rare diseases, it is often necessary to draw case-families from multiple racial/ethnic groups. We provide a summary of our recommendations in **Table 1**.

### Study limitations

We only consider methods from an extensive literature search with open-source implementation and an available manual. The focus of this work is on a dichotomous trait and common markers. We have not considered methods for quantitative traits [Laird and Lange, 2008], or methods tailored to rare variants, which are increasingly becoming available via high-throughput whole exome or whole genome sequencing techniques [Hecker et al., 2020]. We do not assess any indirect effects (e.g., maternal effect, parent-of-origin effect, imprinting) or any interaction effect (e.g., maternal-fetal genotype interactions, gene-environment interaction, epistasis) [Ainsworth et al., 2011]. We use the TDT-like methods exclusively as association tests [Laird and Lange, 2006], and do not explore their type I error rates separately for the other possible null hypothesis scenarios under the original “no linkage *or* no association” composite null hypothesis [Laird and Lange, 2008, Hecker, Laird and Lange, 2019]. We do not consider monads [Fan et al., 2013] or any other pedigree structure [Chen, Manichaikul and Rich, 2009, Hecker, Laird and Lange, 2019]. We explore three different mother-child pair designs and their corresponding trio designs; other hybrid designs are certainly possible [Vermeulen et al., 2009, Gjerdevik et al., 2020] but are beyond the scope of this paper. We consider bi-allelic SNPs only (not considering multi-allelic or haplotype effects) [Cordell, Barratt and Clayton, 2004, Gjessing and Lie, 2006]. Our simulation framework is simple and do not reflect the usual complex genetic architecture underlying many disorders; however, we use a similar framework and parameter choices as used by many others [Deng and Chen, 2001, Chen, Manichaikul and Rich, 2009, Hecker, Laird and Lange, 2019]. Our simulations do not involve any confounders since most TDT-like methods cannot accommodate covariate effects. We focus on either one or two homogenous genetic ancestry groups to consider effects of population substructure but do not consider the full range of admixture.

Nonetheless, it is important to bear in mind that we have undertaken the first attempt at benchmarking these popular methods across dyad and trio designs, across a multitude of modeling approaches and under different data types and structure. We provide some practical guidelines for an appropriate selection of methods to use in different potential scenarios present in consortium studies such as ECHO, or any other nuclear family-based study designs.

## Software

gTDT: https://www.bioconductor.org/packages/release/bioc/html/trio.html

TDT, 1-TDT, GDT-PO: https://www.chen.kingrelatedness.com/software/GDT/index.shtml

TDT_D+C_, TDT_DC_ : https://github.com/RayDebashree/TDT-like-tests

HAPLIN: https://cran.r-project.org/web/packages/Haplin/index.html

## Data Availability Statement

The GENEVA data on clefts are publicly available on dbGaP (https://www.ncbi.nlm.nih.gov/gap/, study accession number phs000094.v1.p1).

## Supplemental Data

The online supplementary materials provide additional figures.

## Acknowledgments

This research was supported in part by the NIH for the Environmental influences of Child Health Outcomes Data Analysis Center (U24OD023382), and the R03DE029254 (DR, THB). All analyses were carried out using computing cluster—the Joint High Performance Computing Exchange—at the Department of Biostatistics, Johns Hopkins Bloomberg School of Public Health. We thank Dr. Wei-Min Chen for providing us the LE program for simulating pedigree data, Dr. Min Shi for pointing us to relevant literature on log-linear models in dyad designs, and Dr. Ingo Ruczinski for helpful discussions relating to family-based designs in general.

