## Supplemental Data for "Benchmarking statistical methods for analyzing parent-child dyads in genetic association studies"

**Figure S1:** Type I error performance of the different methods at stringent levels for a combined analysis of dyads and trios when the distribution of dyads and trios are skewed across ancestry groups. Results are based on 1 million null SNPs simulated on two genetic ancestry groups, where 500 case-mother dyads come from one group, and 250 case-mother dyads plus 250 case-parent trios come from the other. The two ancestry groups are assumed to have common disease prevalence of 30% and 15% respectively, and the causal SNP has MAF 10% and 3% respectively. All offspring are affected, and all parents are unaffected. Observed($-\log_{10}$p-values) are plotted on the y-axis and Expected($-\log_{10}$p-values) on the x-axis of the QQ plots. The gray shaded region in each QQ plot represents a conservative 95% confidence interval for the expected distribution of p-values.

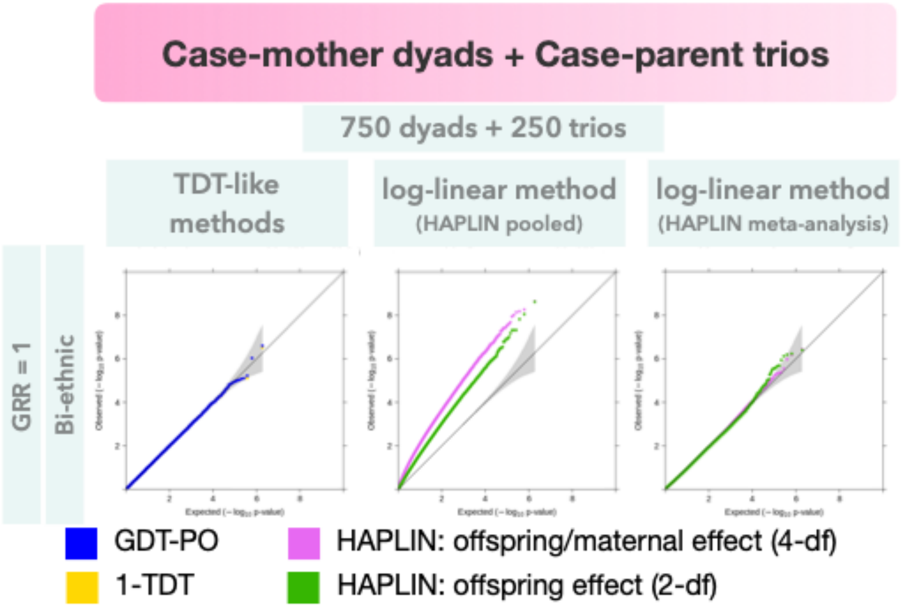

**Figure S2:** Type I error performance of the different combinations of hybrid designs at stringent significance levels when the distribution of case and control families are skewed across ancestry groups. Results are based on 1 million null SNPs simulated on two genetic ancestry groups, where 500 case families (case-parent trios or case-mother dyads as the case may be) come from one group, and 500 control families come from the other. The two ancestry groups are assumed to have common disease prevalence of 30% and 15% respectively, and the causal SNP has MAF 10% and 3% respectively. Case-to-control ratio among offspring is 50:50, and all parents are unaffected. Observed($-\log_{10}$p-values) are plotted on the y-axis and Expected($-\log_{10}$p-values) on the x-axis of the QQ plots. The gray shaded region in each QQ plot represents a conservative 95% confidence interval for the expected distribution of p-values.

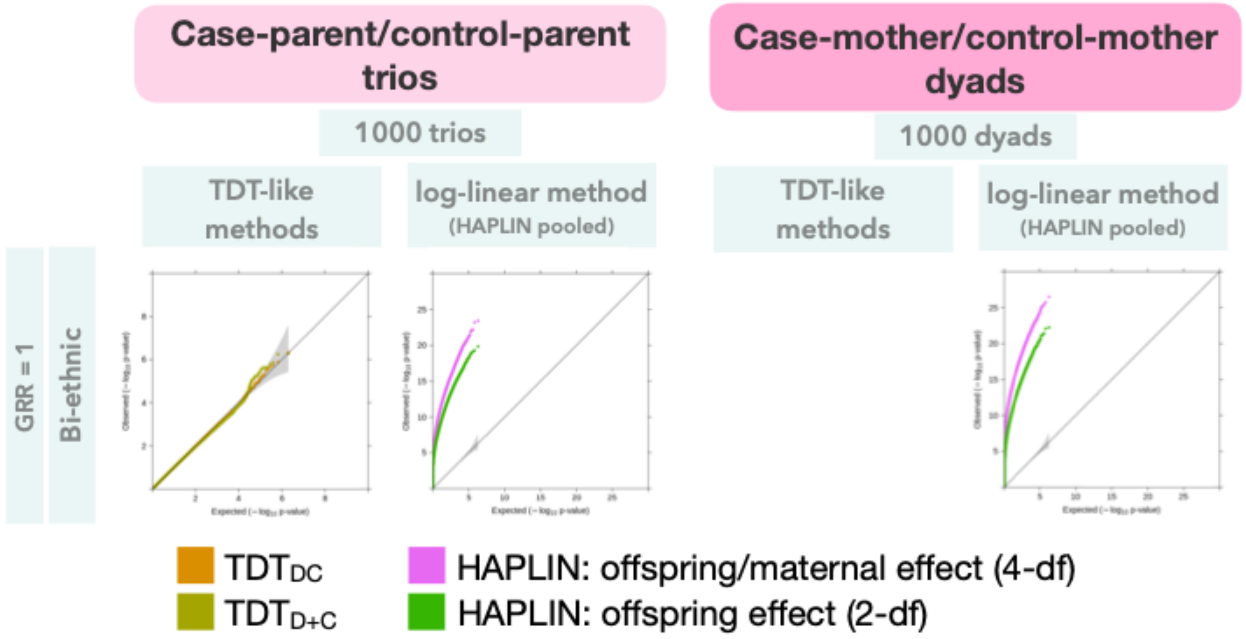

**Figure S3:** Manhattan plots for HAPLIN (2-df test of offspring genotypic effect) when applied on the pooled sample of multi-ethnic families from the GENEVA study on CL/P. The red and blue horizontal lines in each plot correspond to genome-wide ($5 \times 10^{-8}$) and suggestive ($10^{-6}$) significance levels, respectively.

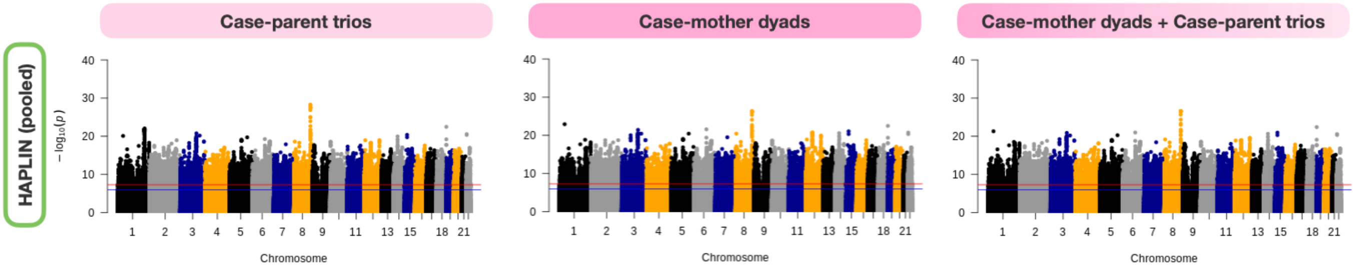
